# Fit by design: Developing substrate-specific seed mixtures for functional dike grasslands

**DOI:** 10.1101/2023.03.09.530576

**Authors:** Markus Bauer, Jakob K. Huber, Johannes Kollmann

## Abstract

1. Sowing is a well-established restoration technique to overcome dispersal limitation. Site-specific seed mixtures are most effective to achieve functional communities. This is especially important if the restored vegetation has to protect critical infrastructure like roadsides and dikes. Here, an improved seed–substrate combination will secure slope stability, reduce mowing efforts, and generate species-rich grasslands.
2. A factorial field experiment addressed this topic on a dike at River Danube in SE Germany in 2018–2021. Within 288 plots, we tested three sand admixtures, two substrate depths, two seed densities and two seed mixture types (mesic hay meadow, semi-dry calcareous grassland) in north and south exposition, and measured the recovery completeness by calculating the successional distance to reference sites, the persistence of sown species, and the Favourable Conservation Status (FCS) of target species.
3. Overall, the sown vegetation developed in the desired direction, but a recovery debt remained after four years, and some plots still showed similarities to negative references from ruderal sites. In north exposition, hay meadow-seed mixtures developed closer to the respective reference communities than dry-grassland mixtures.
4. In south exposition, the sown communities developed poorly which might be due to a severe drought during establishment. This initial negative effect remained over the entire observation period.
5. Sand admixture had a slightly positive effect on target variables, while substrate depth, seed density and mixture type had no effects on species persistence or FCS.
6. Synthesis and applications: Site-adapted seed mixtures make restoration more effective. However, applying several seed–substrate combinations might foster beta diversity. Furthermore, additional management efforts are recommended, as they might be necessary to reduce the recovery debt, as well as re-sowing after unfavourable conditions.

## Introduction

Grasslands can support an exceedingly high biodiversity and they provide several ecosystem services (Bardgett et al., 2021; Dengler et al., 2014). However, they are globally endangered (Bardgett et al., 2021), and in Europe, calcareous grasslands and hay meadows are red-listed habitats (Category 3, ‘vulnerable,’ Janssen et al., 2016). Restoration is seen as a key factor to sustain biodiversity and ecosystem services (*Convention on Biological Diversity* (*CBD*), 2014; United Nations, 2019), and sowing is a well-established approach to establish species-rich grasslands (Kiehl et al., 2010). Sowing high-diversity mixtures of local provenance produced by specialized companies is a promising way to scale up restoration efforts (Freitag et al., 2021), and to overcome dispersal filters (Myers & Harms, 2009; Orrock et al., 2023). However, there are still open questions about adjusting seed mixtures to specific site conditions and future climate conditions (Török et al., 2021).

Restoration ecology can increase the predictability of restoration approaches (Mouquet et al., 2015) by using rigorous, repeatable, and transparent experiments based on advanced theory, which will finally strengthen evidence-based restoration (Cooke et al., 2018; Wainwright et al., 2018). Local site conditions and the restoration method are key predictors for vegetation development after sowing (Brudvig et al., 2017), while habitat and biotic filtering are the main assembly factors which can be manipulated by the choice of seed–substrate combinations (Török & Helm, 2017). This means a close adaptation of the substrate to the niche of the target species or of the seed mixtures to the characteristics of the chosen substrate. Suitable substrates reduce habitat filtering of the seeded species, while specific seed mixture minimises competitive exclusion of desired species and simultaneously prohibiting invasive species (Funk et al., 2008). Modifying seed mixtures to match the site conditions could be based on functional plant traits (Balazs et al., 2020; Funk et al., 2008; Laughlin, 2014), although this is not easy to implement (Bauer et al., 2022; Merchant et al., 2022). This challenge is particularly interesting for artificial substrates that are used in urban areas (Bauer et al., 2022), in quarries (Chenot-Lescure et al., 2022), or on dikes (Liebrand & Sykora, 1996).

Dikes are promising sites for the restoration of species-rich grasslands because they can increase habitat area and connectivity of semi-natural grasslands and therefore significantly contribute to biodiversity conservation in agricultural landscapes (Bátori et al., 2020). Steep slopes with different exposition, contrasting substrate layers and dense swards for erosion protection characterise these habitats (Bátori et al., 2016; Berendse et al., 2015; Husicka, 2003). Dikes can reconcile several ecosystem functions including both flood security and rich biodiversity (Teixeira et al., 2022), which can be fostered by an adapted seed–substrate combination.

The aim of this study is to identify the best combinations of seed mixtures and substrates for vital and species-rich grasslands on north- and south-exposed dike slopes. An experiment was set up to test different substrate depths, sand admixtures, seed densities, and mixture types. We expected a better development of dry grassland in the south exposition with shallow and sandy substrates, and of mesic meadows in north exposition on less sandy and deeper substrates. For steep slopes, e.g., on dikes, high seed densities are recommended for successful vegetation establishment (Kleber-Lerchbaumer et al., 2017), albeit without experimental evidence.

The success of restoration, i.e., the difference from desired conditions, is evaluated by comparing the species composition with reference sites (cf. Brudvig et al., 2017), since the successional distance to reference grasslands describes the recovery completeness (Rydgren et al., 2019). Furthermore, we observed the persistence, which is the presence of the sown species monitored over three consecutive years (Wilsey, 2021). Finally, the Favourable Conservation Status (FCS) was calculated which distinguishes habitat-characteristic diversity and non-typical derived diversity (Helm et al., 2015). Based on four years of monitoring, we tested the following hypotheses:

1. Site conditions on northern vs southern dike slopes facilitate establishment of mesic or dry grassland mixtures, respectively.
2. Nutrient reduction by sand addition and shallow substrates increase the establishment of dry-grassland compared to mesic seed mixtures.
3. Reduced soil resources benefit target species of species-rich grasslands.
4. High seed densities increase the establishment of sown plants and suppress non-target species.

## Materials and methods

### Field experimental design

Specific combinations of seed mixtures and substrates (‘seed–substrate combinations’) were tested on a dike at the Danube River in SE Germany (Figure **1**; 314 m a.s.l.; WGS84: lat/lon, 48.83895/12.88412). The climate of the region is temperate-suboceanic with a mean annual temperature of 8.4 °C and an annual precipitation of 984 mm (Deutscher Wetterdienst, 2021). During the study, three exceptionally dry years (2018–2020) occurred (Appendix A1, Hari et al., 2020), as well as three minor floods, which, though, did not reach the plots (Appendix A1). The substrates consisted of calcareous sand (0–4 mm) and agricultural soil obtained from a nearby dike construction site near the village of Steinkirchen. A big roller mixed both components and an excavator put the substrates in the prepared plots.

**Figure 1:**
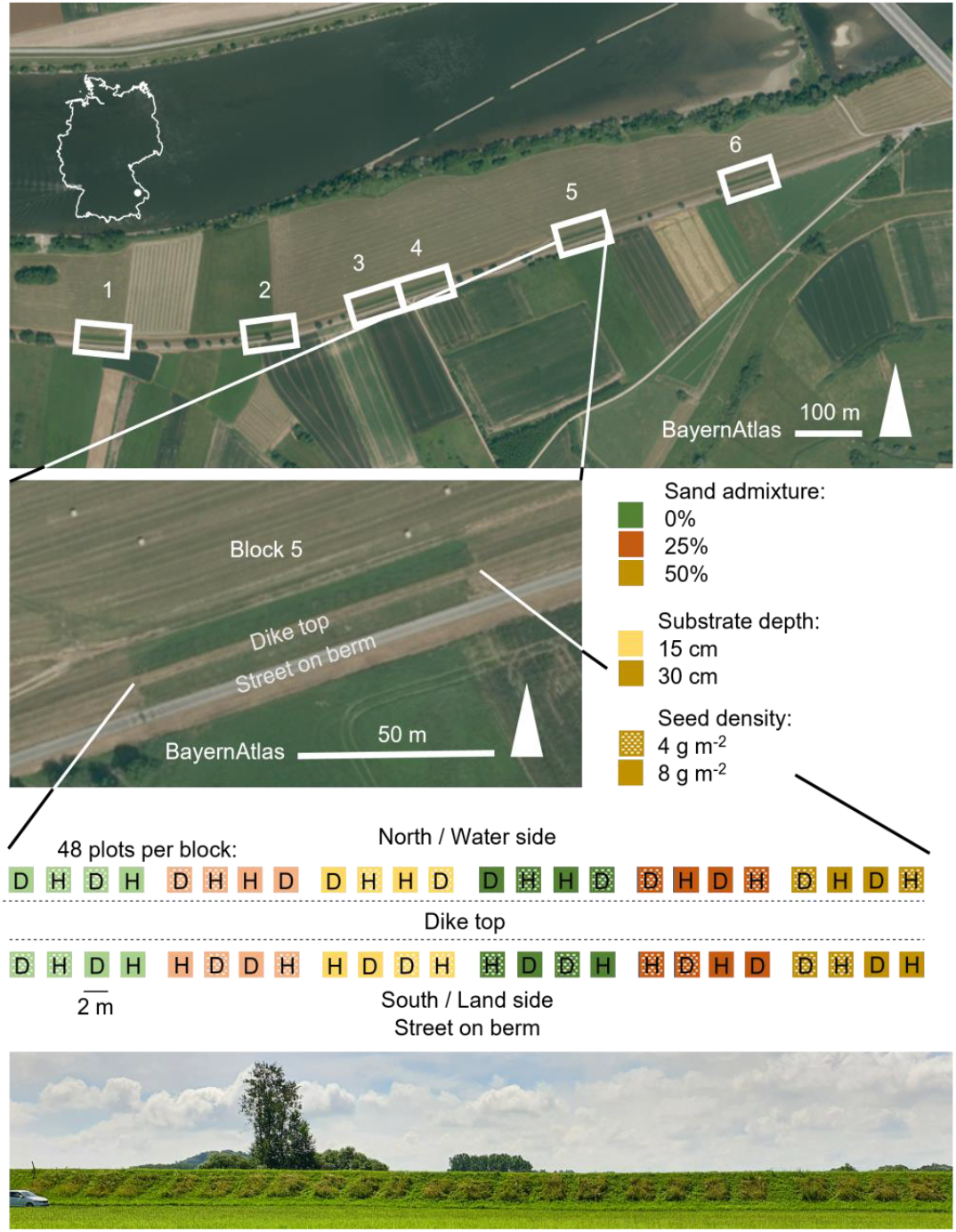
Local setting and design of the multifactorial experiment on grassland sowing on dikes. The experiment was located on a dike at River Danube in SE Germany. The 288 plots were allocated in six blocks (white squares on the upper photo) and on the north and south slope (central photo) (both aerial photos: Bayerische Vermessungsverwaltung, 2023). Four treatments were conducted: sand admixture, substrate depth, seed density, and seed mixture types H and D (hay meadows, dry grasslands). The western half of a block had a shallow substrate depth and within this, half of the substrates had different sand admixtures. The photo on the bottom shows the northern slope of one block in 2021, four years after sowing (photo: Markus Bauer).

The target vegetation types were lowland mesic hay meadows and semi-dry calcareous grassland (EUNIS codes: R22, R1A, Chytrý et al., 2020; Arrhenatherion elatioris and Cirsio-Brachypodion pinnati according to the EuroVegChecklist: CM01A, DA01B, Mucina et al., 2016). The species pool of hay meadows and dry grasslands consisted of 55 and 58 species, respectively. The seeds were received from a commercial producer of autochthonous seeds (Co. Krimmer, Pulling). From these species pools, 20 species were selected for each plot in a stratified, randomised manner (Appendix A2). Each mixture contained seven grasses (60wt% of total seed mixture), three legumes (5%) and ten herbs (35%) (Table **1**). The hay-meadow mixtures had higher community-weighted means (CWM) for specific leaf area (SLA), lower for seed mass, and higher for canopy height than the dry-grassland mixtures (Appendix A3). The south-exposed plots were sown in mid-April 2018 and the north exposition 14 days later. In October 2018, *Bromus hordeaceus* was sown as a nursery grass to provide safe sites under drought conditions. In late-April 2018 due to the drought, the south exposition was protected by a geotextile consisting of straw chaff (350 g m^−2^) which was removed after two weeks due to unsatisfactory effects on seedling emergence. The management started with a cut at 20 cm height without hay removal in August 2018, followed by standard deep cuts with hay removal in July 2019 and 2020.

**Table 1.** Each plot received an individual set of twenty species with some restrictions to the number of species per functional group. The total species pool for hay meadows was 55 and for dry grassland 58. All individual seed mixtures are stored in Appendix A2.

| Functional group | Species pool |  | Seed mixture | Total ratio | Ratio per species |
| --- | --- | --- | --- | --- | --- |
|  | Hay meadow | Dry grassland |  |  |  |
|  | # | # | # | wt% | wt% |
| High grasses | 6 | 5 | 3 | 25.7 | 8.6 |
| Low grasses | 8 | 8 | 4 | 34.3 | 8.6 |
| Legumes | 5 | 7 | 3 | 5.0 | 1.7 |
| Herbs | 34 | 36 | 9 | 30.0 | 3.3 |
| Hemiparasites | 2 | 2 | 1 | 5.0 | 5.0 |

We used 288 plots of the size 2.0 m × 3.0 m, vertically oriented, halfway up the dike slopes, distributed over the north and south exposition, and arranged in six blocks (=replicates). The experiment used a split-plot design combined with a randomised complete block design (Figure **1**). The split plot was created by the two expositions of the dike, where all 24 treatment combinations were tested, i.e., sand admixtures (0, 25, and 50%), soil depths (15 vs. 30 cm), the two seed mixture types, and two seed densities (4 vs. 8 g m^−2^, cf. Kiehl et al., 2010; Kleber-Lerchbaumer et al., 2017).

Below the substrate, a 5-cm thick drainage layer of gravel (0–16 mm) was installed. Soil samples of the three substrates from both expositions were tested by mixing several sub-samples from different plots. The sand admixture changed the soil texture, increased the C/N ratio, reduced calcium carbonate, but did hardly change the pH which was within the weak alkaline range (Table **2**). The pH and C/N ratio were within the recommended range, as well as the clay ratio of the 25% sand admixture and the substrate depth of 30 cm (Husicka, 2003). Phosphate and potassium were rather scarce for agricultural soils, but magnesium showed high concentrations (Bayerisches Landesamt für Landwirtschaft (LfL), 2022).

**Table 2.** Characteristics of the substrates used for the sowing experiment on river dikes. Soil samples of the three substrates were analysed for the fraction <2 mm. The soil texture was classified according to the ‘Bodenkundliche Kartieranleitung’ (Bundesanstalt für Geowissenschaften und Rohstoffe, 2005). The pH was measured in CaCl_2_ solution. Plant available phosphorus and potassium were measured in a calcium acetate-lactate extract and magnesium in a CaCl_2_ extract. For calculating CaCO_3_, a sub-sample was annealed at 550 °C and the measured C amount multiplied with 8.33. To calculate total N and the C/N ratio, a sub-sample was incinerated at 1000 °C. Lt3 = medium clayey loam; Ls4 = strong sandy loam; Sl3 = medium loamy sand; Sl4 = strong loamy sand.

| Exposition | Sand admixture | Skeleton (>2 mm) | Sand | Silt | Clay | Soil texture | pH | N | P<sub>2</sub>O<sub>5</sub> | K<sub>2</sub>O | Mg<sup>2+</sup> | C/N | CaCO<sub>3</sub> |
| --- | --- | --- | --- | --- | --- | --- | --- | --- | --- | --- | --- | --- | --- |
|  | vol% | vol% | wt% | wt% | wt% |  |  | wt% | mg 100 g<sup>-1</sup> | mg 100 g<sup>-1</sup> | mg 100 g<sup>-1</sup> |  | wt% |
| North | 0 | 5 | 18 | 45 | 37 | Lt3 | 7.4 | 0.35 | 4 | 6 | 27 | 8.9 | 12.1 |
|  | 25 | 26 | 49 | 29 | 22 | Ls4 | 7.4 | 0.24 | 4 | 5 | 25 | 9.0 | 8.8 |
|  | 50 | 40 | 75 | 14 | 11 | Sl3 | 7.5 | 0.11 | 3 | 4 | 17 | 9.5 | 5.3 |
| South | 0 | 9 | 18 | 45 | 37 | Lt3 | 7.3 | 0.37 | 6 | 7 | 28 | 8.8 | 12.5 |
|  | 25 | 26 | 59 | 23 | 18 | Ls4 | 7.4 | 0.19 | 3 | 5 | 23 | 9.2 | 7.3 |
|  | 50 | 44 | 71 | 18 | 13 | Sl4 | 7.5 | 0.13 | 4 | 5 | 16 | 9.5 | 7.3 |

### Vegetation surveys

The vegetation was surveyed in June or July 2018–2021 (Braun-Blanquet, 1964) and the Londo scale was used (Londo, 1976). The establishment rates of species were recorded in Appendix A4. Establishment success was high with 48 species of the species pool of hay meadows (87%) and 46 (79%) of dry grasslands recorded by 2021, which are rather good ratios (cf. Hedberg & Kotowski, 2010); the species established in 31 ± 22% (mean ± SD) of their sown plots. In total, 274 vascular plant species were found (Appendix A5).

The recovery completeness was described by the successional distance which quantifies the distance of a plot to the average reference site in the ordination (*d_jt,0_*, Rydgren et al., 2019, Figure **2**). Persistence was derived from the ‘species losses’ component of the temporal beta-diversity index (TBI; 1 – *B*_sor_) which was calculated by comparing the seed mixtures with the respective species composition of each year using Sørensen dissimilarity (Legendre, 2019). The Favourable Conservation Status (FCS) is the ratio of characteristic and derived diversity measured as species richness (Helm et al., 2015). Characteristic diversity consists of species that belong to a habitat-specific species pool and derived diversity consists of all other species. The habitat-specific species pool consisted of all sown species and other typical species of mesic and dry grasslands (Appendix A5).

**Figure 2:**
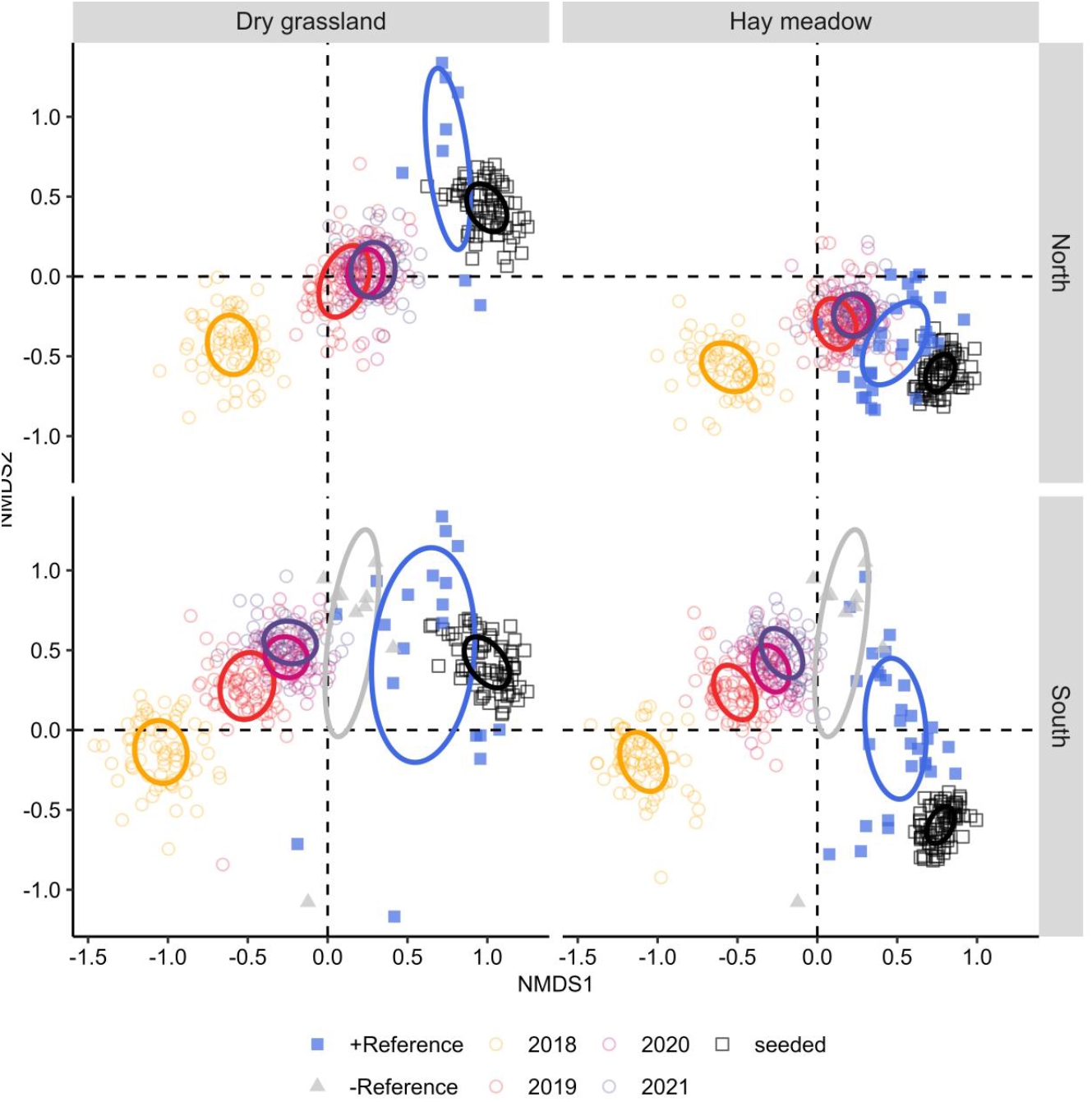
The species composition of sown experimental plots on a river dike over time and in comparison with reference sites and the seed mixtures. Both expositions and both seed mixture types are shown in separate panels. The NMDS was based on the Sørensen dissimilarity and data of 288 plots observed over four years after sowing in 2018 (circles). These experimental plots were compared with the seed mixtures (black squares) and 98 positive and negative reference plots (filled symbols) from older dike grasslands in the surroundings (Bauer et al., 2023a), and six plots from sPlotOpen (Sabatini et al., 2021). The ellipses show the standard error of the groups. 2D-stress: 0.21.

To compare the restoration outcomes with real references and not solely with seed mixtures, vegetation surveys were extracted from sPlotOpen (Sabatini et al., 2021) and our own surveys on the Danube dikes in the surroundings (Bauer et al., 2023a). We selected six dry grassland plots (EUNIS code R1A, Chytrý et al., 2020) within SE Germany from sPlotOpen and 98 plots of our own survey, which included also hay meadows (R22), and as a negative reference ruderal, dry and anthropogenic vegetation (V38).

### Statistical analysis

A non-metric multidimensional scaling ordination (NMDS) with Sørensen dissimilarity was used to visualise variation in species composition in space and time. Seven species were excluded because they had an accumulated cover over all plots of<0.5%. Finally, 343 species were included in the ordination.

To measure the effects of the treatments on our three response variables, we calculated Bayesian linear mixed-effects models (BLMM) with the random effect plot nested in block with the Cauchy prior (see Lemoine, 2019). Furthermore, we included as a fixed effect the botanists, who recorded a certain plot. For the simple effects of the treatments, we chose plausible weakly informative priors. To evaluate the influence of the priors, prior predictive checks and models with non-informative priors were calculated.

For the computation, we used four chains, a thinning rate of two, 5,000 iterations for warm-up, and 10,000 in total. We used the Markov Chain Monte Carlo method (MCMC) with the no-U-turn Sampler (NUTS). For evaluating the computation, the convergence of the four chains was checked using trace plots and evaluating *R*-hat values, and MCMC chain resolution by the effective sampling size (ESS). Posterior predictive checks were done with Kernel density estimates histograms of statistics skew and leave-one-out (LOO) cross-validation (see Gabry et al., 2019). Finally, the models were compared with the Bayes factor (BF) and Bayesian *R*^2^ values (Gelman et al., 2019).

Data, code and the entire model specifications and evaluations are stored on GitHub and presented in an easily accessible document for scrolling through (Bauer et al., 2023b). There, the sections are referenced to the Bayesian analysis reporting guidelines (BARG, Kruschke, 2021). All analyses were performed in R (Version 4.2.2, R Core Team, 2022), with the functions ‘brm’ from the package ‘brms’ (Bürkner, 2017) for model calculation, several functions from ‘brms’ and ‘bayesplot’ (Gabry & Mahr, 2022) for model evaluation, and ‘metaMDS’ from ‘vegan’ for the ordination (Oksanen et al., 2022).

## Results

### Hay meadows on north exposition closer to reference

The ordination showed the species composition of seed mixtures and the development of the plots during four years (Figure **2**). The NMDS confirmed that the seed mixtures were variable, albeit distinctive for hay meadows and dry grasslands and confirming the intended direction of the vegetation development. As one exception, hay meadows in south exposition did not develop towards their seed-mixture compositions.

The reference sites had a larger variation than the seed mixtures and were close to the seed mixtures but hardly overlapped (Figure **2**). The positions of the reference sites shifted to the left in comparison to the seed mixtures, which means in the direction of early-successional stages. Nonetheless, they still differed from the negative references of ruderal vegetation. Negative references were only available on the south exposition and they were located in the NMDS between the positive reference sites and the state of restored plots in 2021. Nevertheless, 33% of the 288 plots reached the state of the target habitat types by 2021 (EUNIS code R22, R1A, Chytrý et al., 2020). Hay meadow-seed mixtures led to a closer development to hay-meadow references than dry grasslands to their references (Figure **3**A, **4**A). This was especially the case in north exposition (Figure **2**).

**Figure 3:**
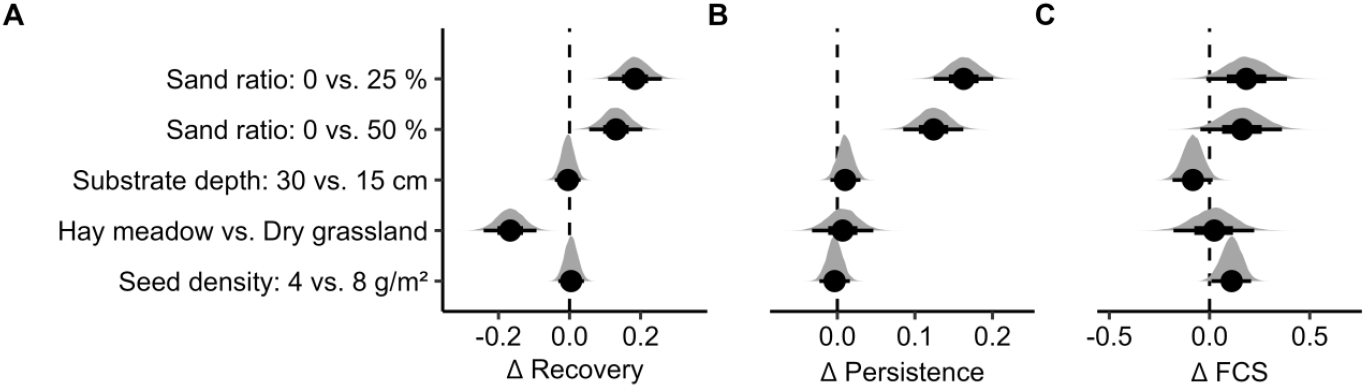
Effects of treatments on the development of sown grassland communities at a river dike. The posterior density distributions (grey) are calculated over all four surveyed years and both expositions. Shown are the medians, 66% and 95% credible intervals, which were derived from a Bayesian linear mixed-effects model (BLMM). Shown are (A) the recovery completeness compared to reference sites, (B) the persistence of sown species, and (C) the Favourable Conservation Status (FCS). The FCS is the ratio of target species to non-target species. Note that the zero lines indicate that both levels have equal values. This means, e.g., that hay meadows are closer to their reference than dry grasslands (A).

**Figure 4:**
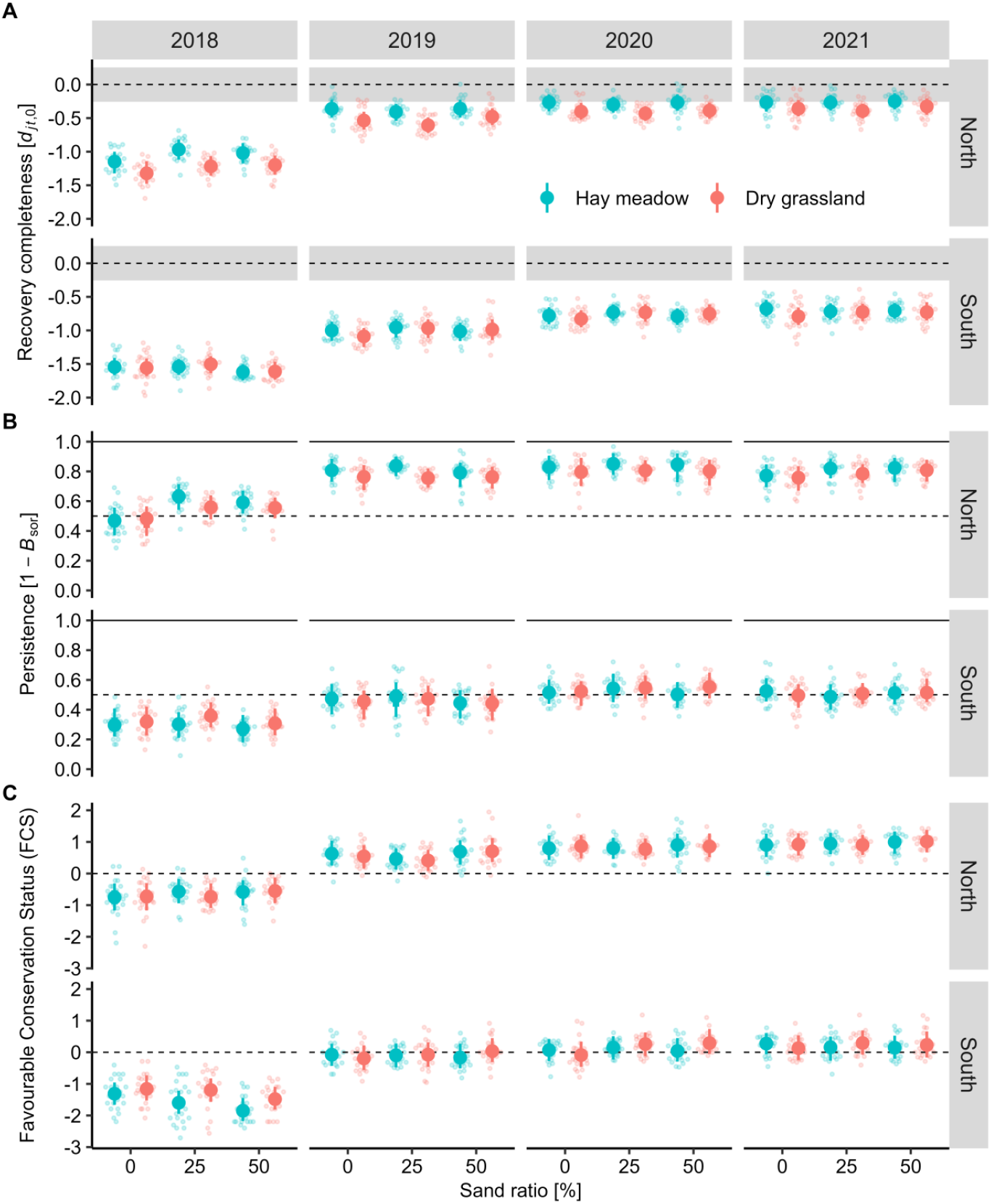
The development of grassland communities at a river dike over four years after sowing. The plots had substrates with different sand admixtures and were sown with two different seed mixture types. Three indices are evaluated. (A) Recovery completeness (*d_jt,0_*): the zero lines indicate the mean position of the reference sites for each habitat type on the NMDS axis 1 (Figure **2**). The grey area marks the standard deviation of the position of the reference sites (Figure **2**). (B) Persistence of sown species: losses component of the temporal beta-diversity index (1 – *B*_sor_). (C) Favourable Conservation Status (FCS): the zero line indicates that target and non-target species are balanced. Positive values indicate that there are more target species. Shown are the medians and 95% credible intervals of the posterior distributions, which were derived from a Bayesian linear mixed-effects model (BLMM).

### Weak effects of substrates and seed density

We could identify a statistically clear positive effect of the sand admixture on the persistence of sown species and on the recovery rate, but no effects of seed density or substrate depth (Figure **3**). The posterior distributions are also shown in the interaction plots that separate exposition and survey year (Figures **4**). For all three response variables, the vegetation developed positively after one year, while the recovery rate slowed down in the following years. Both expositions revealed similar trends but for all responses, the values were clearly lower in south exposition, e.g., persistence values were on average more than 46% higher in north exposition (Figure **4**B). The interactions of restoration treatments were neither clear nor strong. Persistence of both seed mixture types was slightly positively affected by sand admixture in north exposition (+ 6–7 ± 4%, Figure **4**B).

## Discussion

### Success of the restoration approaches

The seed mixtures and their positive reference sites were similar but hardly overlapped (Figure **2**). The position on the ordination suggests that the seed mixture represents a late-successional stage compared to the references. The NMDS shows a slightly better adaptation of hay meadows to the north exposition than of dry grasslands (Figure **4**A). This can be expected from the requirements of hay meadows for mesic conditions, which can be provided on north-exposed dike slopes (Bátori et al., 2020; Oberdorfer, 1993). In southern exposition, the plots of hay meadows developed rather towards dry grassland references which indicates an ineffective restoration due to a non-adapted seed mixture.

The vegetation developed generally in the desired direction but was still distinct from positive references and seed mixtures after four years. In the south exposition, the plots were rather similar to the negative reference of dry ruderal vegetation. The gap between goal and restoration outcome was also shown for other sowing experiments or restorations (Engst et al., 2016; Kaulfuß et al., 2022; Mitchley et al., 2012) or for dike vegetation compared with semi-natural reference grassland (Bátori et al., 2016). This result is not surprising since the ‘recovery debt’ is a general phenomenon of grassland restoration (Jones et al., 2018; Moreno-Mateos et al., 2017).

### General effects of treatments and exposition

Restoration on agricultural soils can have limited success due to high nutrient loads (Walker et al., 2004) but mixing with a mineral component need not necessarily improve the outcome (Chenot-Lescure et al., 2022). Similarly to a study in France, sand admixture reduced nutrient loads and led to higher persistence of sown species while a 50% admixture did not further increase this effect. In addition, the effect only appeared in north exposition and the effect size of about 6% in the 4^th^ year of restoration was rather small. The Favourable Conservation Status (FCS) was hardly affected by the sand admixture, which corresponds to an experiment in a quarry (Chenot-Lescure et al., 2022). Substrate depth did not significantly affect persistence or FCS, similarly to earlier studies (Baer et al., 2004; Husicka, 2003). Larger differences in soil depths might be necessary to observe negative effects by thicker substrate layers as was shown for prairies (Dornbush & Wilsey, 2010) or a substrate depth of<15 cm, since most roots occur in the topsoil on dikes (Vannoppen et al., 2016). Seed density had also no clear effect on persistence and FCS which fits the results of Kaulfuß et al. (2022), who found that a certain amount of seeds is necessary for a successful establishment of target species, but higher densities do not further improve the outcome, but rather have a slightly negative effect.

The vegetation in south exposition had a different species composition, which confirms the findings of Bátori et al. (2016) in Hungary. However, the differences might also be due to methodical reasons, since the geotextile, which had been implemented on the southern slope, was removed after two weeks. This was unfortunate for at least some seedlings, and amplified by the intense drought in summer 2018 and 2019 (cf. Hari et al., 2020; Larson et al., 2021; Orrock et al., 2023). The lasting negative effect on persistence and FCS on the southern slope suggests a legacy effect of adverse weather conditions after sowing as observed by other studies (Groves et al., 2020; Stuble et al., 2017). These conditions during the establishment phase might have led to a special trajectory (Suding et al., 2004), and probably levelled the distinction of the seed mixture types in south exposition.

### No interaction effect of seed–substrate combinations

Our aim was to identify perfect seed–substrate combinations regarding restoration effectiveness and biodiversity. For evaluating effectiveness, we measured the persistence of the sown species, and FCS for investigating plant biodiversity. However, we could not identify an interaction effect for neither one of these indices. We would have expected a better performance of hay-meadow seed mixtures with lower sand admixture and for dry grasslands with higher sand admixture. Our results suggest that, at least after four years, the substrate conditions are within the range of both seed mixture types (hay meadows vs dry grasslands). Although, both types are clearly phytosociologically and functionally distinct, they are still relatively close, because they contain shared species and develop under similar site conditions with modified sub-associations (Appendix A3, Husicka, 2003; Oberdorfer, 1993). Other grassland studies could identify more or less clear interactions of opposing habitat preferences or functional traits along the gradients of productivity, moisture and nutrients (Freitag et al., 2021; Kaulfuß et al., 2022; Zirbel & Brudvig, 2020). However, these studies did not work with an experimental set up of different seed– substrate combinations, but analysed the result of habitat and biotic filtering after 1, 5 and 15yrs, respectively. Furthermore, the non-existence of ideal combinations could be explained by priority effects that means that the species of the imperfect-adapted seed mixture type could establish earlier and pre-empted the available niches for the species of related habitat types (Fukami, 2015).

## Conclusions

Our results suggest that adapted seed mixtures can increase restoration effectiveness by sowing hay meadows in the north but not necessarily in south exposition of dikes. Furthermore, the reduction of the nutrient load through sand admixture was positive, albeit with small effect size. The question remains if sand admixture is the most efficient restoration measure to promote diversity on dikes. Increasing seed density on dike slopes does not appear to be necessary, and soil depths of 30 cm are not adverse compared to 15 cm thick substrates.

There were no perfect seed–substrate combinations and thus we conclude that a variation of seed mixture types and different substrates along restoration sections would promote biodiversity more than a single uniform solution (Bauer et al., 2023a; Holl et al., 2022). Negative effects of drought in the sowing season might require re-sowing. To close the recovery debt, the management adaptation might be promising since this is a crucial factor beside the restoration approach and the site characteristics (Grman et al., 2013; Tölgyesi et al., 2021). For example, the introduction of sheep grazing on the experimental plots, which already exists in the surroundings, will modify the disturbance regime and improve dispersal. Overall, our results support the finding that restored dike grasslands can promote biodiversity in agricultural landscapes (Bátori et al., 2020). However, the recovery debt highlights the fact that restored grasslands cannot substitute old-growth grasslands (Nerlekar & Veldman, 2020).

## Acknowledgements

We would like to thank our project partners Dr. Markus Fischer, Frank Schuster, and Christoph Schwahn (WIGES GmbH) as well as Klaus Rachl and Stefan Radlmair (Government of Lower Bavaria) for numerous discussions on restoration and management of dike grasslands. Fieldwork was supported by Clemens Berger and Uwe Kleber-Lerchbaumer (Wasserwirtschaftsamt Deggendorf). We thank Holger Paetsch, Simon Reith, Anna Ritter, Jakob Strak, Leonardo H. Teixeira, and Linda Weggler for assisting with the field surveys or soil analyses in 2018–2020. The German Federal Environmental Foundation (DBU) supported MB with a doctoral scholarship.

## Author contribution

JH and JK designed the experiment. JH did the surveys in the years 2018–2020, and MB in 2019 and 2021. MB did the analyses and wrote the manuscript. JK and JH critically revised the manuscript.

## Open research

Data and code are stored on Zenodo (Bauer et al., 2023b). Model evaluation is stored on GitHub: https://github.com/markus1bauer/2023_danube_dike_experiment/tree/main/markdown

## Session Info

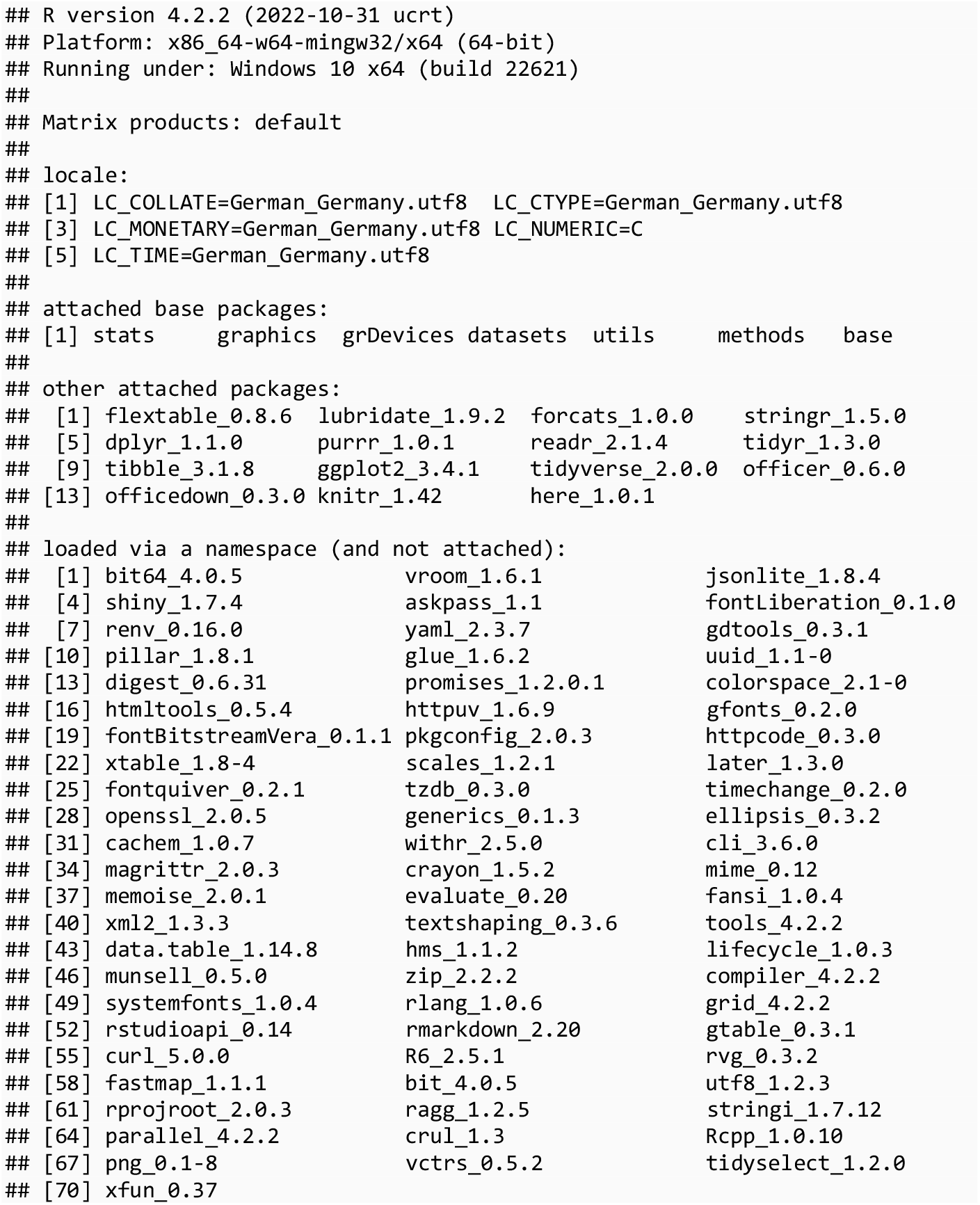

