## Appendix for "Fit by design: Developing substrate-specific seed mixtures for functional dike grasslands"

### Appendix A1

Three dry years (2018–2020) and three minor floods (2018, 2019, 2021) occurred during the study period. Annual temperature, precipitation and discharge of River Danube near the study sites (2017–2021), based on weather station Metten (mean, 1981–2010, ID: 3271, WGS84: lat/lon, 48.85476/12.918911, Climate Data Center of the German Meteorological Service, 2022a, 2022b), stream gauge Pfelling (ID: 10078000, WGS84: lat/lon, 48.87975/12.74716, Bayerisches Landesamt für Umwelt, 2021a), and water level Deggendorf (ID: 10081004, WGS84: lat/lon, 48.82508/12.96229, Bayerisches Landesamt für Umwelt, 2021b). HQ2 = Highest discharge with a probability of occurrence every second year. HSW = Highest water level for shipping.

**Table 1:**

| Environmental variable | Unit | 2017 | 2018 | 2019 | 2020 | 2021 | Long-term average |
|---|---|---|---|---|---|---|---|
| Temperature | °C | 9 | 10.5 | 10.1 | 9.9 | 8.8 | 8.4 |
| Precipitation | mm | 1,014 | 706.0 | 864.0 | 832.0 | 1,034.0 | 984.0 |
| Discharge max (HQ2) | $m^3\ s^{-1}$ | 1,050 | 1,630.0 | 1,440.0 | 1,360.0 | 1,450.0 | 1,620.0 |
| Water level max (HSW) | cm | 567 | 706.0 | 662.0 | 650.0 | 669.0 | 620.0 |

### Appendix A2

#### Generating species pools

For both target vegetation types, a respective initial plant species collection was assembled. The two lists contained the following groups:

- Character species of the corresponding phytosociological units (Arrhenatherion and Mesobromion, as listed in standard literature) and of FFH-habitat types 6510 and 6210 as described for Bavaria
- Typical species of corresponding habitat types according to official directives for biotope mapping in Bavaria
- Species consistently occurring in our own vegetation records on existing river dikes of the region, conducted in 2017, and in species-rich grasslands, recorded within a wide-range habitat documentation project in the Danube valley near Straubing and Deggendorf from 2011. This project had been commissioned as planning grounds for state flood protection programs.

The two species pools were derived from this collection by removing the following groups:

- Very rare and/or regionally absent species
- Woody shrubs, which were excluded from dike vegetation guidelines in Germany because of vortex erosion phenomena around their rooting system
- Nitrogen indicators
- Neophytic and/or invasive species or species with rapid clonal propagation with a high potential of forming mono-dominant stands
- Species of strong ruderal character
- Ephemeral/annual species
- Species, for which no seeds were available in the market or which are difficult to establish by propagule application, e.g., *Carex* spec. or cryptogams
- Taxonomically and functionally closely related and similar species

#### Constructing seed mixtures

For constructing concrete seed mixtures of 20 pant species, a pre-defined number of species within certain functional groups was selected randomly. This way, seed mixtures consisted of three tall grasses, four small grasses, three legumes, one hemiparasite (*Rhinanthus* spec.) and nine herbal perennials, three of which listed as character species in the literature. The actual choice was conducted in a manner, so that each individual species was assigned to each block and each treatment level combination with a near-constant frequency. Furthermore, all species of each functional group were chosen almost the same number of times. In total, plots received 24 or 48 g of seeds (4 vs. 8 g $m^{-2}$).

All sown seed mixtures can be found on GitHub in the folder **‘outputs/tables/ table_a2_seed_mixtures.csv’**. There, we indicate which species were combined and their respective ratio in the specific seed mixture.

### Appendix A3

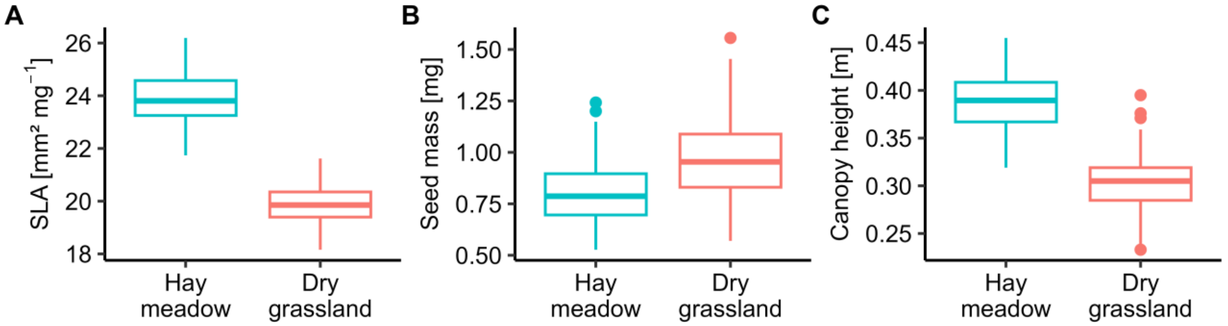


**Figure 1:**

The community-weighted means (CWM) of the seed mixtures used for the dike experiment are based on abundance data: specific leaf area (SLA), seed mass and canopy height. The CWMs were weighted by seed weight in the seed mixture and indicate differences between the target habitat types hay meadows and calcareous grasslands. The functional traits of specific leaf area (SLA), seed mass and canopy height (available traits of all species: 99.2, 98.8, 99.6%) were retrieved from the databases LEDA (Kleyer et al., 2008) and TRY (Kattge et al., 2020).

*Acknowledgment*

This figure was supported by the TRY initiative on plant traits (http://www.try-db.org). The TRY initiative and database is hosted, developed and maintained by J. Kattge and G. Boenisch (Max Planck Institute for Biogeochemistry, Jena, Germany). TRY is currently supported by Future Earth/bioDISCOVERY and the German Centre for Integrative Biodiversity Research (iDiv) Halle-Jena-Leipzig.

### Appendix A4

Establishment rate of sown species (Nomenclature: World Flora Online, 2021). Indicated is the number of sown plots and the percentage of these plots that were occupied by the respective species (presence-absence data). This means that for each species only the sown plots were included. The shades of grey indicate the success of establishment.

**Table 2:**

| Species | Family | Sown plots [#] | Establishment rate [%] | | | |
|---|---|---|---|---|---|---|
| | | | 2018 | 2019 | 2020 | 2021 |
| *Agrostis capillaris* | Poaceae | 160 | 0 | 2 | 1 | 17 |
| *Alopecurus pratensis* | Poaceae | 56 | 0 | 4 | 2 | 2 |
| *Anthoxanthum odoratum* | Poaceae | 158 | 0 | 18 | 33 | 31 |
| *Arrhenatherum elatius* | Poaceae | 144 | 16 | 41 | 46 | 44 |
| *Brachypodium pinnatum* | Poaceae | 74 | 0 | 0 | 9 | 51 |
| *Briza media* | Poaceae | 158 | 0 | 0 | 27 | 46 |
| *Bromus erectus* | Poaceae | 144 | 11 | 40 | 49 | 47 |
| *Cynosurus cristatus* | Poaceae | 45 | 0 | 2 | 13 | 22 |
| *Dactylis glomerata* | Poaceae | 58 | 3 | 43 | 47 | 59 |
| *Festuca ovina* | Poaceae | 73 | 0 | 36 | 34 | 22 |
| *Festuca pratensis* | Poaceae | 130 | 0 | 44 | 46 | 35 |
| *Festuca rubra* | Poaceae | 159 | 1 | 40 | 36 | 29 |
| *Helictotrichon pubescens* | Poaceae | 72 | 0 | 3 | 29 | 43 |
| *Holcus lanatus* | Poaceae | 86 | 0 | 31 | 36 | 29 |
| *Koeleria pyramidata* | Poaceae | 71 | 0 | 1 | 15 | 38 |
| *Phleum pratense* | Poaceae | 128 | 0 | 21 | 11 | 17 |
| *Poa angustifolia* | Poaceae | 74 | 0 | 19 | 23 | 15 |
| *Poa compressa* | Poaceae | 51 | 0 | 37 | 20 | 0 |
| *Poa pratensis* | Poaceae | 47 | 0 | 9 | 11 | 34 |
| *Trisetum flavescens* | Poaceae | 58 | 2 | 45 | 53 | 57 |
| *Anthyllis vulneraria* | Fabaceae | 61 | 33 | 49 | 61 | 33 |
| *Hippocrepis comosa* | Fabaceae | 61 | 2 | 0 | 8 | 2 |
| *Lathyrus pratensis* | Fabaceae | 86 | 1 | 9 | 21 | 36 |
| *Lotus corniculatus* | Fabaceae | 148 | 36 | 49 | 45 | 34 |
| *Medicago falcata* | Fabaceae | 62 | 2 | 35 | 52 | 56 |
| *Medicago lupulina* | Fabaceae | 148 | 9 | 48 | 69 | 13 |
| *Onobrychis viciifolia* | Fabaceae | 61 | 56 | 70 | 52 | 23 |

| Species | Family | Sown plots [#] | Establishment rate [%] | | | |
|---|---|---|---|---|---|---|
| | | | 2018 | 2019 | 2020 | 2021 |
| *Trifolium montanum* | Fabaceae | 63 | 0 | 0 | 0 | 0 |
| *Trifolium pratense* | Fabaceae | 86 | 38 | 43 | 38 | 29 |
| *Vicia cracca* | Fabaceae | 88 | 14 | 17 | 28 | 17 |
| *Achillea millefolium* | Asteraceae | 85 | 34 | 49 | 58 | 48 |
| *Agrimonia eupatoria* | Rosaceae | 36 | 0 | 22 | 31 | 44 |
| *Ajuga reptans* | Lamiaceae | 34 | 3 | 12 | 6 | 12 |
| *Anthericum ramosum* | Asparagaceae | 38 | 0 | 0 | 0 | 0 |
| *Arabis hirsuta* | Brassicaceae | 38 | 0 | 21 | 0 | 0 |
| *Asperula cynanchica* | Rubiaceae | 36 | 0 | 25 | 17 | 6 |
| *Buphthalmum salicifolium* | Asteraceae | 35 | 0 | 31 | 49 | 26 |
| *Campanula glomerata* | Campanulaceae | 71 | 0 | 0 | 0 | 0 |
| *Campanula patula* | Campanulaceae | 32 | 0 | 0 | 50 | 12 |
| *Campanula rapunculoides* | Campanulaceae | 35 | 0 | 3 | 6 | 0 |
| *Campanula rotundifolia* | Campanulaceae | 36 | 0 | 0 | 11 | 0 |
| *Carex flacca* | Cyperaceae | 70 | 0 | 0 | 0 | 0 |
| *Carlina vulgaris* | Asteraceae | 35 | 0 | 0 | 0 | 31 |
| *Centaurea jacea* | Asteraceae | 36 | 44 | 64 | 78 | 53 |
| *Centaurea scabiosa* | Asteraceae | 85 | 25 | 45 | 42 | 36 |
| *Centaurea stoebe* | Asteraceae | 38 | 87 | 92 | 89 | 76 |
| *Cerastium fontanum ssp vulgare* | Caryophyllaceae | 35 | 0 | 54 | 49 | 23 |
| *Cichorium intybus* | Asteraceae | 49 | 47 | 67 | 47 | 49 |
| *Cirsium oleraceum* | Asteraceae | 36 | 0 | 3 | 0 | 8 |
| *Clinopodium vulgare* | Lamiaceae | 35 | 6 | 37 | 40 | 51 |
| *Crepis biennis* | Asteraceae | 50 | 24 | 48 | 48 | 36 |
| *Daucus carota* | Apiaceae | 48 | 42 | 62 | 60 | 40 |
| *Dianthus carthusianorum* | Caryophyllaceae | 38 | 0 | 34 | 63 | 66 |
| *Echium vulgare* | Boraginaceae | 34 | 94 | 97 | 97 | 88 |
| *Filipendula vulgaris* | Rosaceae | 36 | 0 | 6 | 6 | 19 |
| *Galium mollugo* | Rubiaceae | 35 | 40 | 43 | 51 | 54 |
| *Galium verum* | Rubiaceae | 36 | 14 | 42 | 44 | 42 |
| *Geranium pratense* | Geraniaceae | 34 | 0 | 12 | 32 | 59 |
| *Helianthemum nummularium* | Cistaceae | 33 | 0 | 0 | 0 | 0 |
| *Hypericum perforatum* | Hypericaceae | 35 | 0 | 3 | 3 | 29 |
| *Hypochaeris radicata* | Asteraceae | 72 | 44 | 50 | 49 | 31 |
| *Inula salicina* | Asteraceae | 37 | 0 | 0 | 0 | 0 |
| *Knautia arvensis* | Caprifoliaceae | 70 | 13 | 40 | 60 | 61 |

| **Species** | **Family** | **Sown plots [#]** | **Establishment rate [%]** | | | |
|---|---|---|---|---|---|---|
| | | | **2018** | **2019** | **2020** | **2021** |
| *Leontodon hispidus* | Asteraceae | 35 | 9 | 37 | 46 | 29 |
| *Leucanthemum vulgare* | Asteraceae | 73 | 18 | 48 | 62 | 30 |
| *Origanum vulgare* | Lamiaceae | 69 | 0 | 29 | 42 | 57 |
| *Pastinaca sativa* | Apiaceae | 35 | 0 | 60 | 60 | 60 |
| *Peucedanum oreoselinum* | Apiaceae | 34 | 0 | 0 | 0 | 0 |
| *Pimpinella major* | Apiaceae | 34 | 0 | 0 | 3 | 26 |
| *Pimpinella saxifraga* | Apiaceae | 35 | 0 | 0 | 6 | 3 |
| *Plantago lanceolata* | Plantaginaceae | 51 | 45 | 35 | 55 | 63 |
| *Plantago media* | Plantaginaceae | 35 | 0 | 14 | 26 | 17 |
| *Potentilla recta* | Rosaceae | 36 | 0 | 0 | 50 | 50 |
| *Prunella grandiflora* | Lamiaceae | 35 | 0 | 0 | 0 | 6 |
| *Prunella vulgaris* | Lamiaceae | 35 | 0 | 20 | 34 | 49 |
| *Ranunculus acris* | Ranunculaceae | 34 | 0 | 0 | 29 | 24 |
| *Ranunculus bulbosus* | Ranunculaceae | 72 | 1 | 26 | 21 | 0 |
| *Rhinanthus alectorolophus* | Orobanchaceae | 145 | 0 | 41 | 37 | 28 |
| *Rhinanthus minor* | Orobanchaceae | 143 | 0 | 38 | 14 | 0 |
| *Rumex acetosa* | Polygonaceae | 34 | 26 | 53 | 56 | 35 |
| *Salvia pratensis* | Lamiaceae | 82 | 48 | 73 | 77 | 55 |
| *Sanguisorba minor* | Rosaceae | 70 | 43 | 93 | 90 | 86 |
| *Sanguisorba officinalis* | Rosaceae | 51 | 16 | 8 | 14 | 25 |
| *Scabiosa columbaria* | Caprifoliaceae | 36 | 0 | 0 | 17 | 31 |
| *Sedum acre* | Crassulaceae | 36 | 3 | 47 | 58 | 11 |
| *Silaum silaus* | Apiaceae | 36 | 0 | 0 | 11 | 3 |
| *Silene flos-cuculi* | Caryophyllaceae | 33 | 0 | 0 | 9 | 0 |
| *Silene latifolia ssp alba* | Caryophyllaceae | 34 | 35 | 47 | 44 | 44 |
| *Silene vulgaris* | Caryophyllaceae | 36 | 64 | 97 | 94 | 86 |
| *Thymus praecox* | Lamiaceae | 37 | 5 | 35 | 51 | 41 |
| *Tragopogon pratensis* | Asteraceae | 32 | 56 | 78 | 28 | 16 |
| *Verbascum lychnitis* | Scrophulariaceae | 34 | 0 | 35 | 35 | 29 |
| *Veronica chamaedrys* | Plantaginaceae | 45 | 0 | 24 | 29 | 47 |

### Appendix A5

All species which occurred 2018–2021 and the amount of plots in which they appeared (maximum 1152 = 4 years times 288 experimental plots). In total, 274 different species were found. The shades of grey indicate the amount of established plots.

**Table 3:**

| Name | Seeded | Target species | Presence [# plots] |
|---|---|---|---|
| *Acer campestre* | | | 4 |
| *Acer platanoides* | | | 3 |
| *Acer pseudoplatanus* | | | 112 |
| *Achillea millefolium* | 1 | 1 | 399 |
| *Aegopodium podagraria* | | | 28 |
| *Agrimonia eupatoria* | 1 | 1 | 91 |
| *Agrostis capillaris* | 1 | 1 | 213 |
| *Agrostis stolonifera* | | | 49 |
| *Ajuga reptans* | 1 | 1 | 120 |
| *Alchemilla vulgaris* | | 1 | 1 |
| *Allium scorodoprasum* | | 1 | 10 |
| *Alopecurus myosuroides* | | | 13 |
| *Alopecurus pratensis* | 1 | 1 | 61 |
| *Amaranthus retroflexus* | | | 121 |
| *Anagallis arvensis* | | | 4 |
| *Angelica sylvestris* | | | 1 |
| *Anthericum ramosum* | 1 | 1 | 38 |
| *Anthoxanthum odoratum* | 1 | 1 | 354 |
| *Anthyllis vulneraria* | 1 | 1 | 184 |
| *Apera spica-venti* | | | 1 |
| *Arabis hirsuta* | 1 | 1 | 48 |
| *Arenaria serpyllifolia* | | | 322 |
| *Arrhenatherum elatius* | 1 | 1 | 445 |
| *Artemisia vulgaris* | | | 18 |
| *Asperula cynanchica* | 1 | 1 | 54 |
| *Avena fatua* | | | 2 |
| *Barbarea vulgaris* | | | 21 |
| *Bellis perennis* | | | 1 |
| *Brachypodium pinnatum* | 1 | 1 | 142 |

| Name | Seeded | Target species | Presence [# plots] |
|---|---|---|---|
| *Brassica napus* | | | 1 |
| *Brassicaceae* | | | 5 |
| *Briza media* | 1 | 1 | 285 |
| *Bromus erectus* | 1 | 1 | 368 |
| *Bromus hordeaceus* | | 1 | 753 |
| *Bromus inermis* | | | 584 |
| *Bromus sterilis* | | | 6 |
| *Bunias orientalis* | | | 204 |
| *Buphthalmum salicifolium* | 1 | 1 | 75 |
| *Calystegia sepium* | | | 494 |
| *Campanula glomerata* | 1 | 1 | 72 |
| *Campanula patula* | 1 | 1 | 60 |
| *Campanula rapunculoides* | 1 | 1 | 45 |
| *Campanula rotundifolia* | 1 | 1 | 46 |
| *Capsella bursa-pastoris* | | | 284 |
| *Cardamine hirsuta* | | | 1 |
| *Carex acutiformis* | | 1 | 2 |
| *Carex caryophyllea* | | 1 | 15 |
| *Carex filiformis* | | | 6 |
| *Carex flacca* | 1 | 1 | 75 |
| *Carex hirta* | | 1 | 131 |
| *Carex leporina* | | | 2 |
| *Carex muricata* | | 1 | 12 |
| *Carex praecox ssp praecox* | | 1 | 7 |
| *Carex spec* | | | 11 |
| *Carlina vulgaris* | 1 | 1 | 46 |
| *Centaurea jacea* | 1 | 1 | 148 |
| *Centaurea scabiosa* | 1 | 1 | 224 |
| *Centaurea spec* | | | 1 |
| *Centaurea stoebe* | 1 | 1 | 196 |
| *Cerastium arvense* | | 1 | 99 |
| *Cerastium fontanum ssp vulgare* | 1 | 1 | 341 |
| *Cerastium glomeratum* | | | 40 |
| *Chaerophyllum bulbosum* | | | 3 |
| *Chenopodium album* | | | 406 |
| *Chenopodium polyspermum* | | | 141 |
| *Cichorium intybus* | 1 | 1 | 273 |

| Name | Seeded | Target species | Presence [# plots] |
|---|---|---|---|
| *Cirsium arvense* | | | 36 |
| *Cirsium oleraceum* | 1 | 1 | 41 |
| *Cirsium vulgare* | | | 4 |
| *Clinopodium vulgare* | 1 | 1 | 105 |
| *Consolida regalis* | | | 5 |
| *Convolvulus arvensis* | | | 931 |
| *Cornus controversa* | | | 3 |
| *Cota tinctoria* | | | 2 |
| *Crepis biennis* | 1 | 1 | 242 |
| *Crepis capillaris* | | | 1 |
| *Cynosurus cristatus* | 1 | 1 | 75 |
| *Dactylis glomerata* | 1 | 1 | 321 |
| *Daucus carota* | 1 | 1 | 202 |
| *Dianthus carthusianorum* | 1 | 1 | 204 |
| *Dipsacus fullonum* | | | 1 |
| *Echinochloa crus-galli* | | | 166 |
| *Echium vulgare* | 1 | 1 | 409 |
| *Elymus repens* | | | 601 |
| *Epilobium obscurum* | | | 1 |
| *Epilobium parviflorum* | | | 8 |
| *Epilobium spec* | | | 1 |
| *Equisetum arvense* | | | 106 |
| *Eragrostis minor* | | | 1 |
| *Erigeron annuus* | | | 202 |
| *Erigeron canadensis* | | | 52 |
| *Erysimum cheiranthoides* | | | 11 |
| *Euonymus europaeus* | | | 2 |
| *Euphorbia amygdaloides* | | | 1 |
| *Euphorbia cyparissias* | | 1 | 1 |
| *Euphorbia esula* | | | 79 |
| *Euphorbia helioscopia* | | | 104 |
| *Euphorbia verrucosa* | | 1 | 15 |
| *Fallopia convolvulus* | | | 132 |
| *Festuca arundinacea* | | | 16 |
| *Festuca ovina* | 1 | 1 | 176 |
| *Festuca pratensis* | 1 | 1 | 407 |
| *Festuca rubra* | 1 | 1 | 393 |

| Name | Seeded | Target species | Presence [# plots] |
|---|---|---|---|
| *Filipendula ulmaria* | | 1 | 18 |
| *Filipendula vulgaris* | 1 | 1 | 72 |
| *Fragaria vesca* | | | 6 |
| *Fragaria viridis* | | 1 | 21 |
| *Fumaria officinalis* | | | 15 |
| *Galeopsis tetrahit* | | | 24 |
| *Galium aparine* | | | 4 |
| *Galium mollugo* | 1 | 1 | 353 |
| *Galium palustre* | | | 1 |
| *Galium verum* | 1 | 1 | 181 |
| *Geranium dissectum* | | | 32 |
| *Geranium molle* | | | 1 |
| *Geranium pratense* | 1 | 1 | 73 |
| *Geranium pusillum* | | | 1 |
| *Geranium pyrenaicum* | | | 80 |
| *Geum urbanum* | | | 6 |
| *Glechoma hederacea* | | | 101 |
| *Helianthemum nummularium* | 1 | 1 | 34 |
| *Helianthus annuus* | | | 11 |
| *Helianthus tuberosus* | | | 1 |
| *Helictotrichon pubescens* | 1 | 1 | 138 |
| *Hippocrepis comosa* | 1 | 1 | 72 |
| *Holcus lanatus* | 1 | 1 | 203 |
| *Hordeum vulgare* | | | 10 |
| *Hypericum perforatum* | 1 | 1 | 50 |
| *Hypochaeris radicata* | 1 | 1 | 228 |
| *Inula salicina* | 1 | 1 | 37 |
| *Iris pseudacorus* | | | 1 |
| *Iris spec* | | | 6 |
| *Jacobaea vulgaris* | | | 8 |
| *Knautia arvensis* | 1 | 1 | 343 |
| *Koeleria pyramidata* | 1 | 1 | 121 |
| *Lactuca serriola* | | | 31 |
| *Lamium maculatum* | | | 3 |
| *Lamium purpureum* | | | 5 |
| *Lapsana communis* | | | 4 |
| *Lathyrus pratensis* | 1 | 1 | 242 |

| Name | Seeded | Target species | Presence [# plots] |
|---|---|---|---|
| *Lathyrus sylvestris* | | | 1 |
| *Lathyrus tuberosus* | | | 7 |
| *Leontodon hispidus* | 1 | 1 | 92 |
| *Lepidium campestre* | | | 15 |
| *Leucanthemum vulgare* | 1 | 1 | 338 |
| *Linum spec* | | | 1 |
| *Lolium perenne* | | | 55 |
| *Lotus corniculatus* | 1 | 1 | 421 |
| *Lysimachia nummularia* | | 1 | 88 |
| *Lysimachia vulgaris* | | | 27 |
| *Lythrum salicaria* | | | 1 |
| *Malva spec* | | | 1 |
| *Medicago falcata* | 1 | 1 | 326 |
| *Medicago lupulina* | 1 | 1 | 426 |
| *Medicago minima* | | | 13 |
| *Medicago sativa* | | | 213 |
| *Melilotus albus* | | | 1 |
| *Myosotis arvensis* | | | 63 |
| *Nardus stricta* | | | 5 |
| *Oenothera biennis* | | | 19 |
| *Onobrychis viciifolia* | 1 | 1 | 195 |
| *Ononis spinosa* | | | 2 |
| *Origanum vulgare* | 1 | 1 | 195 |
| *Papaver dubium* | | | 2 |
| *Papaver rhoeas* | | | 225 |
| *Pastinaca sativa* | 1 | 1 | 119 |
| *Persicaria amphibia* | | | 125 |
| *Persicaria bistorta* | | | 3 |
| *Persicaria lapathifolia* | | | 1 |
| *Persicaria minor* | | | 1 |
| *Persicaria mitis* | | | 1 |
| *Peucedanum cervaria* | | | 1 |
| *Peucedanum oreoselinum* | 1 | 1 | 34 |
| *Phleum phleoides* | | 1 | 63 |
| *Phleum pratense* | 1 | 1 | 209 |
| *Pimpinella major* | 1 | 1 | 47 |
| *Pimpinella saxifraga* | 1 | 1 | 51 |

| Name | Seeded | Target species | Presence [# plots] |
|---|---|---|---|
| *Plantago lanceolata* | 1 | 1 | 401 |
| *Plantago major* | | | 6 |
| *Plantago major ssp intermedia* | | | 7 |
| *Plantago media* | 1 | 1 | 85 |
| *Poa angustifolia* | 1 | 1 | 166 |
| *Poa compressa* | 1 | 1 | 86 |
| *Poa pratensis* | 1 | 1 | 195 |
| *Poa trivialis* | | | 69 |
| *Poaceae* | | | 5 |
| *Polygonum aviculare* | | | 122 |
| *Populus alba* | | | 3 |
| *Potentilla anserina* | | | 389 |
| *Potentilla erecta* | | 1 | 6 |
| *Potentilla recta* | 1 | 1 | 76 |
| *Potentilla reptans* | | | 230 |
| *Prunella grandiflora* | 1 | 1 | 43 |
| *Prunella vulgaris* | 1 | 1 | 253 |
| *Prunus avium* | | | 13 |
| *Prunus padus* | | | 2 |
| *Quercus robur* | | | 2 |
| *Ranunculus acris* | 1 | 1 | 96 |
| *Ranunculus bulbosus* | 1 | 1 | 132 |
| *Ranunculus polyanthemos* | | 1 | 3 |
| *Ranunculus repens* | | | 180 |
| *Reseda lutea* | | | 130 |
| *Rhinanthus alectorolophus* | 1 | 1 | 532 |
| *Rhinanthus minor* | 1 | 1 | 227 |
| *Rorippa sylvestris* | | | 387 |
| *Rubus caesius* | | | 356 |
| *Rumex acetosa* | 1 | 1 | 273 |
| *Rumex obtusifolius* | | | 81 |
| *Salix purpurea* | | | 7 |
| *Salix spec* | | | 2 |
| *Salvia pratensis* | 1 | 1 | 338 |
| *Sanguisorba minor* | 1 | 1 | 712 |
| *Sanguisorba officinalis* | 1 | 1 | 154 |
| *Saxifraga granulata* | | 1 | 1 |

| Name | Seeded | Target species | Presence [# plots] |
|---|---|---|---|
| *Scabiosa columbaria* | 1 | 1 | 57 |
| *Securigera varia* | | | 514 |
| *Sedum acre* | 1 | 1 | 82 |
| *Sedum maximum* | | | 1 |
| *Sedum sexangulare* | | | 3 |
| *Setaria pumila* | | | 82 |
| *Setaria spec* | | | 16 |
| *Setaria verticillata* | | | 1 |
| *Setaria viridis* | | | 8 |
| *Silaum silaus* | 1 | 1 | 41 |
| *Silene dioica* | | | 2 |
| *Silene flos-cuculi* | 1 | 1 | 39 |
| *Silene latifolia ssp alba* | 1 | 1 | 143 |
| *Silene vulgaris* | 1 | 1 | 794 |
| *Sisymbrium officinale* | | | 1 |
| *Solidago gigantea* | | | 4 |
| *Sonchus arvensis* | | | 16 |
| *Sonchus asper* | | | 133 |
| *Sonchus oleraceus* | | | 9 |
| *Stachys palustris* | | | 13 |
| *Stachys sylvatica* | | | 4 |
| *Stellaria graminea* | | 1 | 99 |
| *Stellaria media* | | | 21 |
| *Symphytum officinale* | | | 139 |
| *Taraxacum campylodes* | | | 190 |
| *Thalictrum minus* | | 1 | 1 |
| *Thlaspi arvense* | | | 67 |
| *Thymus praecox* | 1 | 1 | 102 |
| *Tragopogon pratensis* | 1 | 1 | 111 |
| *Trifolium arvense* | | 1 | 1 |
| *Trifolium campestre* | | 1 | 16 |
| *Trifolium dubium* | | 1 | 19 |
| *Trifolium hybridum* | | | 5 |
| *Trifolium medium* | | | 1 |
| *Trifolium montanum* | 1 | 1 | 64 |
| *Trifolium pratense* | 1 | 1 | 311 |
| *Trifolium repens* | | | 17 |

| Name | Seeded | Target species | Presence [# plots] |
|---|---|---|---|
| *Tripleurospermum maritimum* | | | 1 |
| *Trisetum flavescens* | 1 | 1 | 306 |
| *Triticum spec* | | | 5 |
| *Ulmus laevis* | | | 2 |
| *Urtica dioica* | | | 7 |
| *Valeriana officinalis* | | 1 | 5 |
| *Valerianella spec* | | | 18 |
| *Verbascum densiflorum* | | | 11 |
| *Verbascum lychnitis* | 1 | 1 | 276 |
| *Verbascum spec* | | | 5 |
| *Verbascum thapsus* | | | 13 |
| *Veronica arvensis* | | 1 | 37 |
| *Veronica chamaedrys* | 1 | 1 | 106 |
| *Veronica persica* | | | 2 |
| *Veronica serpyllifolia* | | 1 | 2 |
| *Veronica spec* | | | 2 |
| *Vicia cracca* | 1 | 1 | 213 |
| *Vicia hirsuta* | | | 2 |
| *Vicia sativa ssp nigra* | | 1 | 19 |
| *Vicia sepium* | | 1 | 32 |
| *Vicia tetrasperma* | | | 1 |
| *Viola arvensis* | | | 6 |
| *Vulpia myuros* | | | 14 |
